# Tumor-Specific CD8^+^ T Cells from the Bone Marrow Resist Exhaustion and Exhibit Increased Persistence in Tumor-Bearing Hosts as Compared to Tumor Infiltrating Lymphocytes

**DOI:** 10.1101/2023.08.28.555119

**Authors:** Elizabeth M. Zawidzka, Luca Biavati, Amy Thomas, Claudio Zanettini, Luigi Marchionni, Robert Leone, Ivan Borrello

**Author notes:** Corresponding Author: Ivan Borrello 1 Tampa General Circle Tampa General Hospital Cancer Institute Tampa FL. **Competing interests:** Ivan Borrello is a co-founder of WindMIL Therapeutics and has intellectual property regarding marrow infiltrating lymphocytes (MILs).

## Abstract

Immunotherapy is now an integral aspect of cancer therapy. Strategies employing adoptive cell therapy (ACT) have seen the establishment of chimeric antigen receptor (CAR)-T cells using peripheral blood lymphocytes as well as tumor infiltrating lymphocytes (TILs) with significant clinical results. Despite these successes, the limitations of the current strategies are also emerging and novel approaches are needed. The bone marrow (BM) is an immunological niche that houses T cells with specificity for previously encountered antigens, including tumor-associated antigens from certain solid cancers. This study sought to improve our understanding of tumor-specific BM T cells in the context of solid tumors by comparing them with TILs, and to assess whether there is a rationale for using the BM as a source of T cells for ACT against solid malignancies. Herein, we demonstrate that T cells from the BM appear superior to TILs as a source of cells for cellular therapy. Specifically, they possess a memory-enriched phenotype and exhibit improved effector function, greater persistence within a tumor-bearing host, and the capacity for increased tumor infiltration. Taken together, these data provide a foundation for further exploring the BM as a source of tumor-specific T cells for ACT in solid malignancies.

**Key Messages:** *What is already known on this topic:* TIL therapy shows efficacy but significant limitations. T cell quality is an important determinant of responses to cellular immunotherapy.

*What this study adds:* T cells from the BM appear superior to TILs in phenotype, transcriptional profile, and function. These differences appear driven by tissue (e.g., bone marrow as compared to tumor).

*How this study might affect research, practice or policy:* The BM could serve as an alternative source of cells for adoptive cellular therapy for solid tumors.

## Introduction

Immunotherapy and adoptive cell therapy (ACT) have transformed the treatment paradigms for several advanced cancers^1–4^. Current ACT approaches utilize peripheral blood lymphocytes (PBLs) or tumor-infiltrating lymphocytes (TILs) from surgically excised tumors to create an infusible cell product^4–6^. Despite promising clinical results, limitations exist with both modalities. PBLs have demonstrated efficacy yet limited persistence and minimal success in infiltrating the immunosuppressive microenvironment of solid tumors^7^. TILs must be harvested from large, metastatic tumors, requiring an advanced disease stage at the time of procurement. Moreover, not every tumor specimen possesses sufficient infiltrating lymphocytes to make this approach broadly feasible^8,9^. Lastly, cells derived from an advanced tumor are known to be dysfunctional, and as such, autologous cell therapy late in disease progression decreases the likelihood of a positive clinical outcome.

The bone marrow (BM) is a reservoir for immunologic memory, where both CD4^+^ and CD8^+^ memory T cells are maintained in a quiescent state^10–12. 10,11,12^These memory cells can extravasate and mount subsequent recall responses following antigen challenge^13–15^. Despite its classic characterization as a primary lymphoid organ, the BM can also function as a secondary lymphoid organ (SLO) and prime naïve T cell responses which generate functional immune effectors and memory cells^16–18^. When classic SLOs such as the lymph nodes and spleen are ablated in mice, homing of transferred T cells to the BM is required to induce effective anti-tumor immunity^19^. Redirection of T cells to the BM induces homeostatic expansion and promotes differentiation of memory precursor cells; it also increases T cell resistance to apoptosis, the ability to produce multiple cytokines, and the capacity for effector expansion, all of which favor improved anti-tumor efficacy^20^. Importantly, the BM contains a high proportion of memory T cells specific for previously encountered antigen^21–23^. Antigen-specific CD4^+^ and CD8^+^ T cells have been identified in the BM: following resolution of acute infections^12^, following localized infection^24^, in the context of ongoing chronic infections^25–28^ or autoimmune diseases^29^, and in the context of cancer, including hematologic^30,31^ (e.g. acute myeloid leukemia, multiple myeloma) and solid^16,32,33^ (e.g. breast, pancreatic, melanoma) malignancies. Importantly, BM T cells have been demonstrated to possess *in vitro* and *in vivo* reactivity to autologous, solid tumors^16,32,34^.

Exhaustion is classically understood as a hypofunctional state that T cells acquire when antigen persists instead of being cleared quickly^35^. Tumor-specific T cells within progressing tumors exhibit key exhaustion hallmarks; they express high levels of inhibitory receptors and are impaired in their ability to proliferate and to produce effector cytokines (IL-2, IFNγ, TNFα)^36–38^. Exhaustion is also a critical adaptive T cell response to chronic stimulation and marked heterogeneity exists within this differentiation trajectory ^39^. Accordingly, subsets of CD8^+^ T cells with distinct roles in exhaustion have been identified in murine and human tumor infiltrates.^39^ A self-renewing, less differentiated population of TCF1^+^ CD8^+^ progenitor exhausted T cells (TPEX) can give rise to effector-like exhausted (TEFF) and terminally exhausted (TEXH) T cells. TEFF cells express CX3CR1 and are required to control chronic infection^40^. Despite their restrained functionality, short-lived TEXH cells are cytolytic and critical in mediating anti-viral and anti-tumor immunity ^39,41–43^. Because of their self-sustaining nature, their ability to give rise to cytolytic cells, and the correlation of their presence with a positive response to checkpoint blockade immunotherapy^44^, TPEX could play a key role in improving immunotherapeutic efficacy^45^.

Using the B16.OVA murine melanoma subcutaneous tumor model, our studies compared endogenous as well as clonotypic tumor specific T cells from the BM with those found in the blood, in lymphoid organs, and in the tumor. We show that tumor-specific CD8^+^ T cells in the BM are less exhausted than TILs in phenotype, transcriptional profile, and in function, and confirm that the BM nurtures memory T cells specific for solid tumors^16,17,32,46^. The BM is enriched for memory T cells (CD62L^+^CD44^+^) that are more stem-like (Ly108^+^TCF1^+^) than TILs which, in contrast, express a classic exhaustion signature (TIM3^+^ TOX^+^ PD1^+^). Considering defined exhaustion paradigms, the BM is enriched in progenitor-like and effector-like cells and devoid of terminally exhausted cells as compared to the TUM. Gene set enrichment analysis (GSEA) and assessment of differential gene expression profiles indicate that BM T cells are transcriptionally more memory-like than their TIL counterparts. Upon restimulation *in vitro*, BM T cells exhibit an improved capacity to produce multiple cytokines. *In vivo*, BM T cells are recovered at frequencies greater than exhausted TILs, both in the acute response to vaccine challenge in healthy hosts and following transfer into tumor-challenged hosts. Lastly, in a competitive transfer into tumor-bearing mice, BM T cells persist longer and show a greater degree of tumor infiltration than TILs. In summary, our findings support the hypothesis that the BM functions as a unique niche for T cells, preserving tumor-specific T cells in a memory-like state while maintaining their ability to respond to cognate antigen challenge. These findings have significant implications for the therapeutic use of BM T cells for ACT against solid tumors.

## Results

### Tumor specific T cells from the bone marrow are more stem-like and memory-like than TILs

To determine whether endogenous tumor-specific T cells could be detected in the BM of mice bearing B16.OVA tumor, we used an OVA-specific tetramer (OVA-Tet) (Fig 1A). The BM maintained a stable, tumor-specific CD8^+^ T cell population (2-5% OVA-Tet+) over the course of tumor progression (Fig S1A, S1B). When examined via high-dimensional UMAP analysis fourteen days after tumor injection (D14), both bulk CD8^+^ and Tet^+^ CD8^+^ T cells from the BM clustered separately from TILs, indicating substantial phenotypic differences in T cells derived from these tissues (Fig 1B). Unsupervised clustering revealed that the BM Tet^+^ CD8^+^ T cells are enriched in expression of Ly108, CD44, CD62L, CD127, and TCF1 (clusters 4 and 7, representing about 7% and 45% of Tet^+^ cells, respectively). By contrast, TILs are largely comprised of exhausted cells expressing TIM3, PD1, and TOX (cluster 5) and effector cells (CX3CR1, KLRG1, cluster 6) (Fig 1C, 1D). Furthermore, the presence of CD62L^+^CD44^+^ cells which largely co-express Ly108 and the stemness marker TCF1 distinguished BM T cells from TILs (Fig 1E). Manual gating confirmed that the BM is enriched for CD8^+^ T cells that are CD62L^+^ CD44^+^ and Ly108^+^ TCF1^+^ (Fig 1F-I). Additionally, BM Tet^+^ T cells consistently maintain higher expression of TCF1 and CD62L than TILs throughout tumor growth (Fig 1J-1K). Thus, endogenous, tumor-specific CD8^+^ T cells are found in the BM and possess a more memory-like phenotype than TILs.

**Figure 1:**
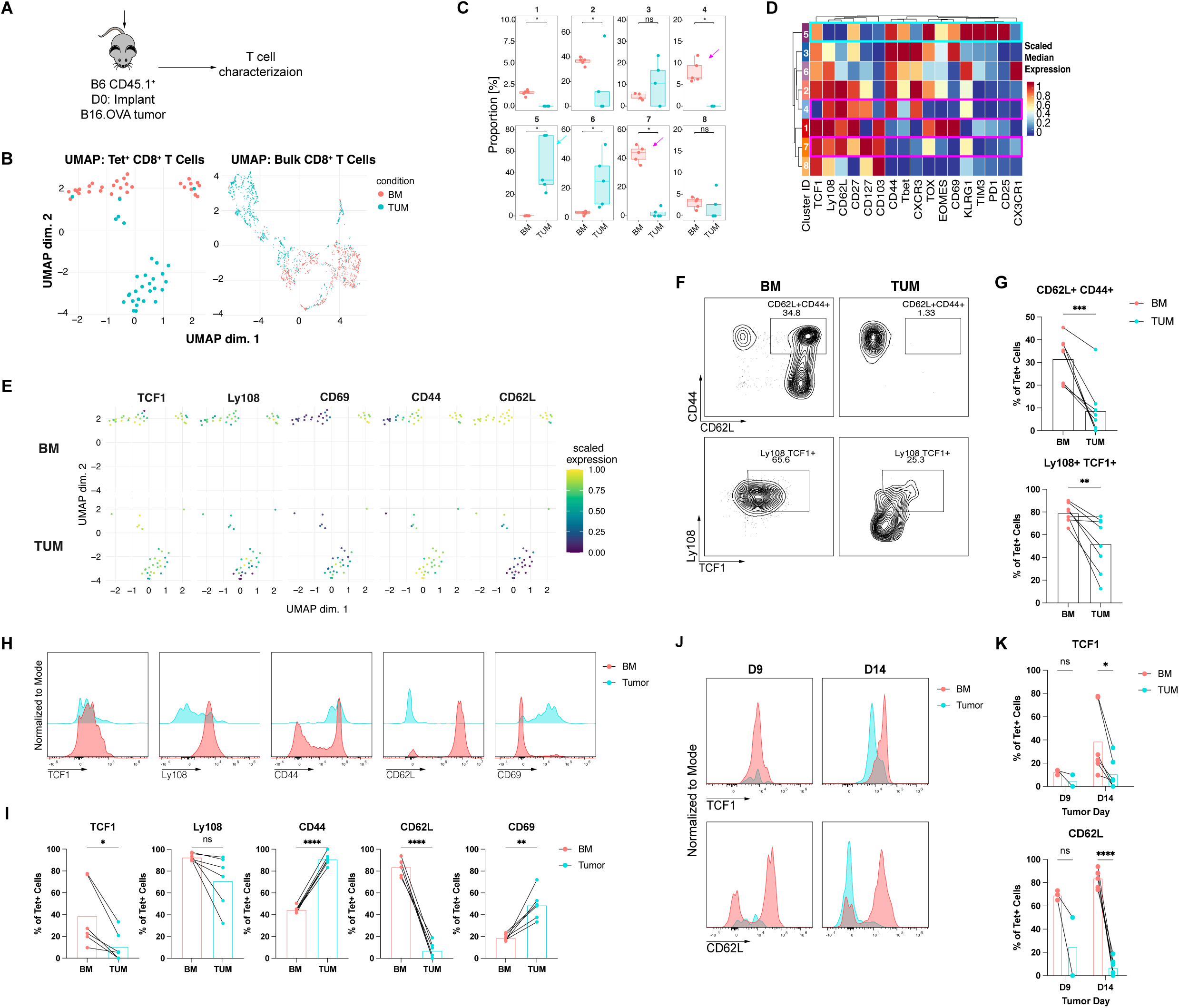
Tumor specific T cells from the bone marrow are more stem-like and memory-like than TILs. **A:** Experimental setup. **B:** UMAP plots of OVA-Tet^+^ CD8^+^ T cells from BM and TUM at tumor D14. **C:** Individual cluster frequencies from unsupervised clustering of data in UMAP analysis. (*, p < 0.05). **D:** Marker expression by individual cluster from UMAP analysis. **E:** UMAP indicating individual marker expression, separated by tissue. **F:** Manual gating on Tet^+^ CD8^+^ T cells from the BM and the TUM. Representative data from two replicate experiments. **G:** Frequencies of CD62L^+^ CD44^+^ and Ly108^+^ TCF1^+^ T cells in the BM and in the TUM at tumor D14. Gated on Tet^+^ CD8^+^ T cells. Data from two replicate experiments analyzed at tumor day 14, n=3-5 mice per group. Analyzed with a paired t test. **, p < 0.01; ***, p < 0.001. **H:** Representative mean fluorescence intensity (MFI) of select markers. Normalized to modal expression. **I:** Quantification of phenotypic marker expression between BM and TUM at B16.OVA D14. Gated on Tet^+^ CD8^+^ T cells and analyzed with a paired t test. *, p < 0.05; **, p < 0.01; ****, p < 0.0001. **J:** MFI indicating changes in expression of TCF1 and CD62L on BM versus TUM samples during tumor progression. Normalized to modal expression. **K:** Frequencies of Tet^+^ cells in BM or in the TUM that are TCF1^+^ or CD62L^+^ over time. Analyzed with multiple paired t tests, *, p < 0.05; ****, p < 0.0001.

### Endogenous, tumor-specific BM T cells express fewer markers of terminal exhaustion than TILs

To contrast BM T cells with TILs, we compared these cell populations based on previously defined exhaustion paradigms. When examined within a framework of progenitor exhausted (Prog1, Prog2, Ly108^+^ CD69^+/-^), intermediate exhausted (TexInt, Ly108^-^ CD69^-^), and terminally exhausted T cells (TexTerm, CD69^+^ Ly108^-^)^39^, the BM is enriched for Tet^+^ cells with a more progenitor-like phenotype. TILs are comprised of all four populations, yet are enriched for TexTerm, consistent with published observations (Fig 2A, 2C). Defining progenitor-like (TPEX) cells as TCF1^+^ TIM3^-^ and terminally exhausted (TEXH) cells as TIM3^+^ TCF1^-47^, BM T cells are enriched for TCF1^+^ TPEX-like cells whereas TILs are skewed toward TIM3^+^ TEXH (Fig 2B, 2C). A recently published schema noted that the valued properties of progenitor-like T cells can be attributed to a Ly108^+^ CD62L^+^ T-SLEX population^43^; in our hands, tumor-specific T cells in the BM are enriched for these markers, whereas this double positive population is notably absent in the tumor (Fig 2B, 2C). Lastly, in a paradigm distinguishing exhausted T cells using Ly108 and CX3CR1 expression (Ly108^+^ TPEX, CX3CR1^+^ effectors (TEFF), and Ly108^-^ CX3CR1^-^ terminally exhausted cells (TTERM))^40^, the BM is enriched for TPEX and TEFF and notably devoid of TTERM cells, in direct contrast to the tumor (Fig 2B, 2C). Thus, regardless of the parameters used to define exhausted T cell subsets, the BM is enriched for CD8^+^ T cells with a progenitor-like phenotype and lacks T cells that appear exhausted.

**Figure 2:**
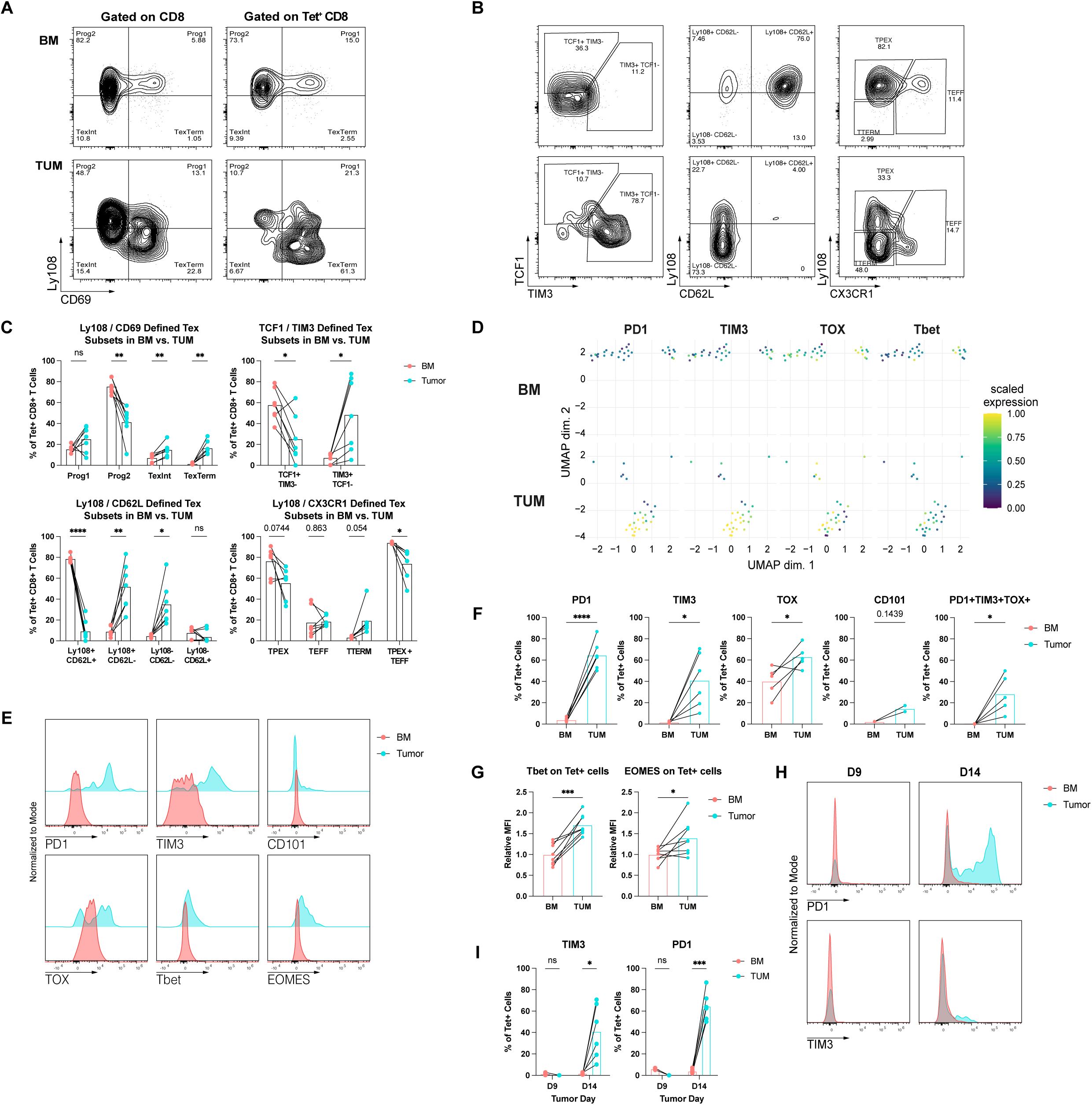
Endogenous, tumor-specific T cells in the bone marrow express fewer markers of terminal exhaustion than TILs. **A:** Endogenous Tet^+^ CD8^+^ T cells from the BM or the TUM gated using exhaustion schema from *Beltra et al, 2020*, in which Ly108 and CD69 are used to define progenitor-exhausted T cells (Prog1, Prog2), intermediate exhausted T cells (TexInt), and terminally exhausted T cells (TexTerm). Representative data from two replicate experiments, tumor D14. **B:** Endogenous BM T cells compared to TILs with three separate gating strategies used to define exhausted T cell subsets in the literature. TCF1^+^ TIM3^-^ progenitor exhausted TPEX or TIM3^+^ TCF1^-^ exhausted TEXH from *Miller et al*, 2019, Ly108^+^ CD62L^+^ TPEX, Ly108^+^ CD62L^-^ TPEX, Ly108^-^CD62L^-^ TEX from *Tsui et al*, 2022, Ly108^+^ TPEX, CX3CR1^+^ TEFF, Ly108^-^ CX3CR1^-^ TEXH from *Zander et al, 2019.* Gated on endogenous Tet^+^ CD8^+^ T cells. **C:** Frequencies of subsets indicated in A-B from BM and TUM at B16.OVA tumor day 14. Gated on endogenous, Tet^+^ CD8^+^ T cells. Data from two replicate experiments, n=3- 5 mice per group. Analyzed with multiple paired t tests, *, p < 0.05; **, p < 0.01; ***, p < 0.001; ****, p < 0.0001. **D:** Individual exhaustion marker expression overlaid on UMAP analysis of Tet^+^ CD8^+^ T cells from the BM or the TUM, separated by tissue. **E:** Relative MFI of exhaustion marker expression on T cells from representative samples in the BM and the TUM. Representative data, gated on Tet^+^ CD8^+^ T cells and normalized to modal expression. **F:** Quantification of exhaustion marker expression between BM and TUM at B16.OVA D14. Gated on Tet^+^ CD8^+^ T cells and analyzed with a paired t test. *, p < 0.05; **, p < 0.01; ***, p < 0.001; ****, p < 0.0001. **G:** Quantification of differences in relative MFI of exhaustion-associated markers between the BM and the TUM at B16.OVA D14. Relative MFI was calculated by normalizing to the average expression of a given marker on BM samples per experiment. Analyzed with a paired t test. *, p < 0.05; **, p < 0.01; ***, p < 0.001; ****, p < 0.0001. **H:** Representative MFI indicating changes in expression of PD1 and TIM3 on BM versus TUM samples during tumor progression. Normalized to modal expression. **I:** Quantification of PD1 or TIM3 expression on Tet^+^ cells from the BM and TUM over time. Analyzed with a paired t test. *, p < 0.05; **, p < 0.01; ***, p < 0.001; ****, p < 0.0001.

Examining individual marker expression computationally, we observed a prominent PD1^+^ TOX^+^ TIM3^+^ population of Tet^+^ cells is present in the TUM and not in the BM (Fig 2D). By mean fluorescence intensity (MFI) as well as with manual gating of key markers on D14, TILs appear more exhausted, further validating these computational findings (Fig 2E-G). Additionally, expression of PD1 and TIM3 increased in the TUM and not in the BM with tumor development (Fig 2H-I). These data indicate that, unlike TILs, endogenous tumor-specific BM T cells are enriched in progenitor-like phenotypes and lack markers of terminal differentiation or dysfunction.

### The BM microenvironment nurtures memory-like, tumor-specific T cells following adoptive transfer

The data presented thus far focus on the endogenous T cell repertoires in the BM and in the TUM. We sought to contrast the fates of antigen-specific T cells and understand their biology within these two compartments. As such, naïve T cells from the spleens of OT-I mice (CD8^+^ T cells expressing receptors with specificity for the SIINFEKL peptide) were transferred into B16.OVA-bearing mice (Fig 3A, Fig S2A-G).

**Figure 3:**
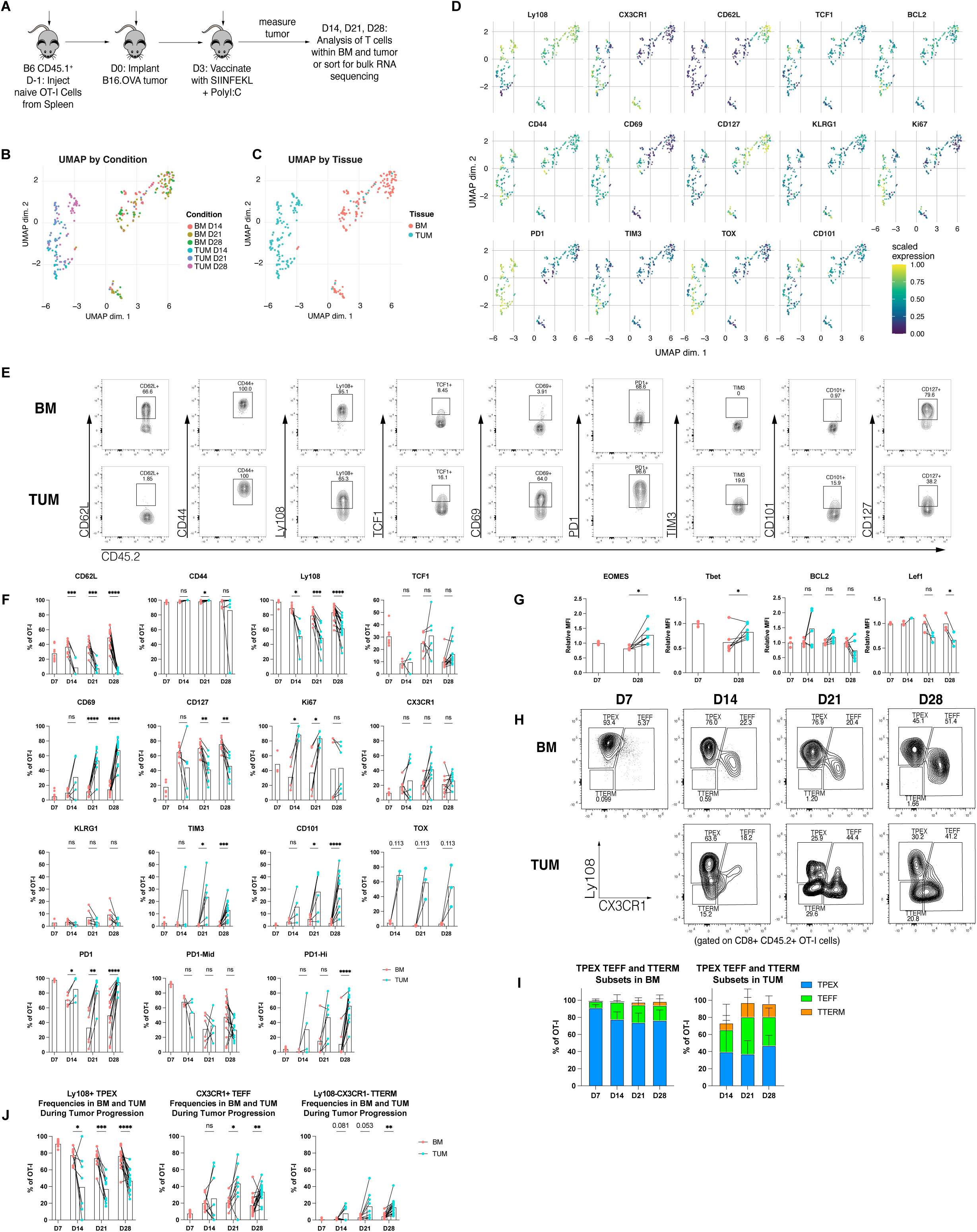
The bone marrow microenvironment nurtures memory-like, tumor-specific T cells following adoptive transfer. **A:** Experimental workflow. **B:** UMAP analysis of OT-I cells from BM and TUM D14, 21, and 28 in adapted model workflow, colored by tissue and timepoint. n=3 mice per timepoint in these analyzed data. **C:** UMAP analysis colored by tissue from which the cells were derived. **D:** Individual marker expression overlaid on UMAP analysis of BM and TUM OT-I cells. **E:** Representative gating of OT-I cells from BM and TUM for select phenotypic markers used to assess differences between these compartments. Gated on CD45.2^+^ CD8^+^ T cells. **F:** Quantification of the expression of surface markers, chemokine receptors, and transcription factors used to assess differences between BM and TUM at different timepoints (B16.OVA D7, D14, D21, and D28) in the new model workflow. Analyzed with multiple paired t tests. *, p < 0.05; **, p < 0.01; ***, p < 0.001; ****, p < 0.0001. **G:** Quantification of the differences in relative MFI of select markers between BM and TUM over the course of tumor development. Relative MFI was calculated by normalizing to the average expression of a given marker on BM samples per experiment. Analyzed with a paired t test. *, p < 0.05; **, p < 0.01; ***, p < 0.001; ****, p < 0.0001. **H:** Timecourse analysis with B16.OVA-OT-I model showing distribution of TPEX, TEFF, TTERM subsets defined with Ly108/CX3CR1 surface markers in the BM or the TUM. Representative data from 4 experiments, n=3-7 mice per group. **I:** Relative proportions of TPEX, TEFF or TTERM cells as percentage of all OT-I cells in the BM or TUM. **J:** Quantification of Ly108^+^ TPEX, CX3CR1^+^ TEFF, and Ly108^-^CX3CR1^-^ TTERM subsets in BM and TUM over the course of tumor development. n=3-7 mice per timepoint, summary of 4 replicate experiments. Analyzed with multiple paired t tests, *, p < 0.05; **, p < 0.01; ***, p < 0.001; ****, p < 0.0001.

Time course data revealed a marked phenotypic concordance between the clonotypic OT-I cells and the endogenous Tet^+^ T cells. Examination of OT-I cells in the BM and TUM demonstrated changes over time. Importantly, examining the same timepoint underscores differences in OT-I cells driven by the tissue from which they are derived (Fig 3B, S3A-C, S4A-D). Consistent with our observations in endogenous T cells, the BM is enriched for CD44^+^ CD62L^+^ memory T cells; BM T cells are largely Ly108^+^ (Fig 3D, Fig S3B), with notable TCF1 and CD127 expression (Fig 3D, Fig S4B, cluster 1). TILs distinguish themselves from BM with a lack of CD62L as well as with greater expression of TIM3, PD1, TOX, and CD101 (Fig 3D-F, Fig S3B, Fig S4B-D, clusters 4- 5). At all timepoints, the BM is enriched for T cells that are CD44^+^CD62L^+^Ly108^+^ (Fig 3F, Fig S4B-D, cluster 2). Cells in the BM upregulate CD127 over time (consistent with a memory phenotype), express Lef1 and TCF1 (indicative of a restrained effector state), express the anti-apoptotic marker BCL2, and cycle less than TILs as measured by Ki67 (Fig 3E-G, Fig S4B-D). Notably, although some BM T cells express PD1, the intermediate expression level suggests activation following antigen encounter. In contrast, the higher PD1 expression in TILs that increases with tumor progression is consistent with a more profound state of T cell exhaustion (Fig 3F). The different trends in key marker expression become more prominent over time, indicating that the BM nurtures different types of cells during tumor progression as compared to the TUM microenvironment.

CX3CR1^+^ CD8^+^ T cells (TEFF) are essential in anti-tumor responses and are distinct from Ly108^+^ TPEX^40^. We used these markers were used to stratify T cell subsets during tumor progression (Fig 3H). The BM maintains a TPEX population of OT-I cells and develops a TEFF population with tumor progression. In contrast, tumor-derived OT-I cells distribute more evenly between TPEX, TEFF, and TTERM populations. Concordant with our prior analyses, the BM is devoid of TTERM cells and significantly enriched for TPEX cells when compared to the tumor where the relative TPEX/TTERM ratio decreases with disease progression (Fig 3H-3J).

Lastly, multidimensional scaling (MDS) analysis of TPEX, TEFF and TTERM subsets at D28 from BM and TUM shows substantial differences between these populations, largely driven by the tissue from which they were acquired (Fig S3C). In fact, BM-and TUM-derived cells separate completely via MDS, independent of TPEX, TEFF or TTERM subset classification. All three marker-defined subsets from the TUM were not statistically distinct in this analysis, while the separation between BM TPEX and TEFF cells indicates distinct phenotypic profiles. This demonstrates that the differences in T cell states are not exclusively driven by subset-defining surface markers (e.g., Ly108 and CX3CR1). Considering that these cells were originally naïve, monoclonal, transgenic OT-I cells, these tissue-driven differences illustrate that the BM and TUM microenvironments enrich for T cells with distinct phenotypes and that the BM is a microenvironment that preferentially nurtures memory-like T cells with progenitor characteristics, even in the context of solid tumor progression.

### Tumor specific BM T cells exhibit a more memory-like and less exhausted transcriptional profile than TILs

To further expand our understanding of the differences in T cell states derived from BM and TUM, OT-I cells from the following subsets were submitted for bulk RNA-sequencing: BM D7 TPEX, TEFF; BM D28 TPEX, TEFF; TUM: D28 TPEX, TEFF, TTERM (Fig 4A). *K means* clustering of the top 500 differentially expressed genes (DEGs) at D28 demonstrates that BM T cells, regardless of subset, are strikingly different from TILs on a transcriptional level (Fig 4B). Likewise, principal component analysis (PCA) of the top 100 DEGs indicates that, despite these T cell subsets being defined by distinct cell surface markers, the predominant distinguishing factor at a transcriptional level is tissue, representing 77% of the variance between samples (PC1). The second principal component (PC2) was T cell subset difference, which only accounted for 6.48% of the variance (Fig 4D).

**Figure 4:**
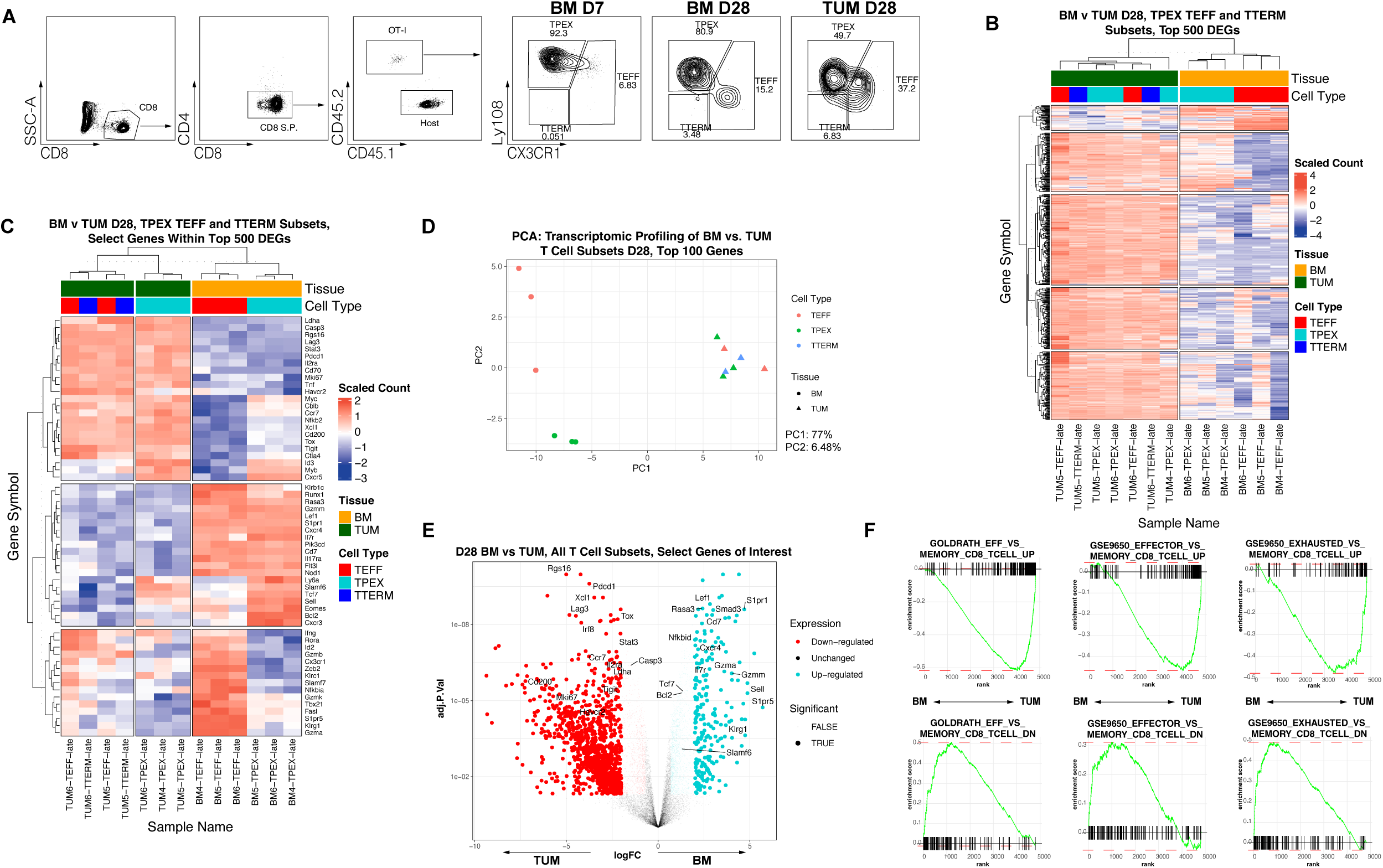
Tumor-specific T cells in the bone marrow exhibit a more memory-like and less exhausted transcriptional profile than TILs. **A:** Gating strategy for bulk RNAseq experiment. **B:** Top 500 differentially expressed genes in T cell subsets derived from BM or TUM at D28, clustered using *k*-means. **C:** Select genes highlighted within top 500 DEGs in T cell subsets from BM or TUM at D28. **D:** Principal component analysis (PCA) of top 100 differentially expressed genes from TPEX, TEFF and TTERM subsets from BM and TUM at D28 of tumor progression. **E:** Volcano plot of BM versus TUM T cells, with differential abundance analysis encompassing all subsets. Select genes of interest highlighted. Upregulated = upregulated in the BM; downregulated = downregulated in the BM and enriched in the tumor. **F:** GSEA of signatures of effector vs. memory CD8^+^ T cells (left and middle) or exhausted vs. memory CD8^+^ T cells (right) in the ranked list of genes differentially expressed by BM-derived T cell subsets versus T cell subsets derived from B16.OVA tumor. Contrast performed using Limma-voom. Individual gene sets met the following criteria: absolute value of the normalized enrichment score ≥ 2 and adjusted p-value < 0.01.

Comparing all BM-derived versus all TUM-derived subsets at D28, BM T cells exhibit enrichment in transcripts associated with memory (*Il7r, Sell, Cd7*), migration (*S1pr1, S1pr5, Cxcr4, Rasa3*), T cell identity and differentiation (*Tcf7*, *Lef1, Runx1, Nod1, Pi3kcd, Flt3l*), and an effector-like state (*Gzmm, Klrb1c, GzmA, Klrg1*). By contrast, TILs express transcripts indicative of a restrained or dysfunctional T cell state (*Pdcd1, Tox, Havcr2, Tigit, Lag3, Cd200, Rgs16*) and an active immune response (*Stat3, Ldha, Tnf, Casp3, Nfkb2, Xcl1, Mki67*) (Fig 4C, 4E). GSEA comparing BM and TUM OT-I cells indicates that BM T cells are significantly enriched for memory-like (versus effector) signatures, as well as for memory-like (versus exhausted) signatures (Fig 4F).

Directly comparing TPEX and TEFF subsets from the BM and the TUM, BM TPEX cells are enriched for a more progenitor-like and memory-like transcriptional profile, as evidenced by significant transcript abundance of *Sell, Lef1, Il7r, Tcf7, Bcl2* (Fig 4C, Fig S5C). BM TEFF cells exhibit a more effector-like transcriptional profile, including enrichment for *Gzma, Gzmm, Gzmk, Klrb1c, Klrg1, S1pr1, Fasl, Rasa3, Zeb2* (Fig 4C, Fig S5D). By contrast, both TPEX and TEFF subsets from the tumor exhibited transcriptional profiles reminiscent of exhaustion, with an abundance of transcripts including *Pdcd1, Tox, Lag3, Havcr2, Tigit, Ctla4, Rgs16* when compared to their analogs in the BM (Fig 4C, Fig S5C-D). These data confirm that tissue microenvironment is a major determinant of transcriptional differences.

BM T cells undergo marked transcriptional changes during tumor progression. In this PCA analysis, PC2 accounted for 22.1% of the variance, delineating differences between D7 and D28 BM T cells (Fig S5A-B). Analysis of BM T cell subsets indicates that BM TPEX and TEFF cells are more similar to each other at D7 than at D28 (Fig S6A-B), suggesting that these subsets take on more distinct roles with tumor progression. At D28, all BM T cells are enriched in transcripts associated with a more quiescent, memory-like state (*Il7r, Bcl2*), whereas at D7 they appear more activated, cycling, and effector-like (*Pdcd1, Gzma, Gzmb, Mki67, Mybl2, Cdk1, Klrg1*) (Fig S6C). GSEA was consistent with these findings (Fig S6E). This suggests that in our model, naïve OT-I cells are activated by vaccine and then take on a more quiescent, memory-like state following arrival in the BM.

Examining subset differences between TPEX and TEFF cells in the BM, D28 TPEX cells in the BM express transcripts associated with a bona fide progenitor-like state (*Tcf1, Xcl1, Ccr7, Id3, Myb, Slamf6*) ^43^, whereas their effector-like counterparts express transcripts associated with an effector state (*Cx3cr1, Gzmb, Gzma, Gzmk, Ifng, Fasl, Klrg1*) and migration (*S1pr5*) (Fig 4C, Fig S6D). GSEA validated that BM TPEX cells are more memory-like and naïve-like than TEFF cells, suggesting a less differentiated state (Fig S6F). These data are consistent with the finding that Ly108^+^ TPEX cells and CX3CR1^+^ TEFF cells possess fundamental biologic differences. Moreover, taken in conjunction with our phenotyping data, these data indicate that the BM is capable of nurturing both progenitor-like and effector-like cells that are tumor specific in the setting of tumor progression.

### BM T cells exhibit greater persistence and tumor infiltration than TILs following adoptive transfer

To assess their capacity for effector cytokine production, we re-stimulated cells from the BM and TUM *in vitro*. Upon restimulation with PMA and ionomycin, T cells from the BM express higher levels of CD107a and can produce higher levels of IFNγ than TILs (Fig 5A, Fig S7A). Additionally, degranulating CD107a^+^ CD8^+^ T cells from the BM exhibit greater polyfunctional (IFNγ^+^ IL-2^+^ TNFα^+^) cytokine production than TILs (Fig 5B, Fig S7B). This indicates that, despite their more quiescent, memory-like state, when maximally stimulated with PMA and ionomycin, BM T cells can simultaneously produce more effector cytokines than TILs.

**Figure 5:**
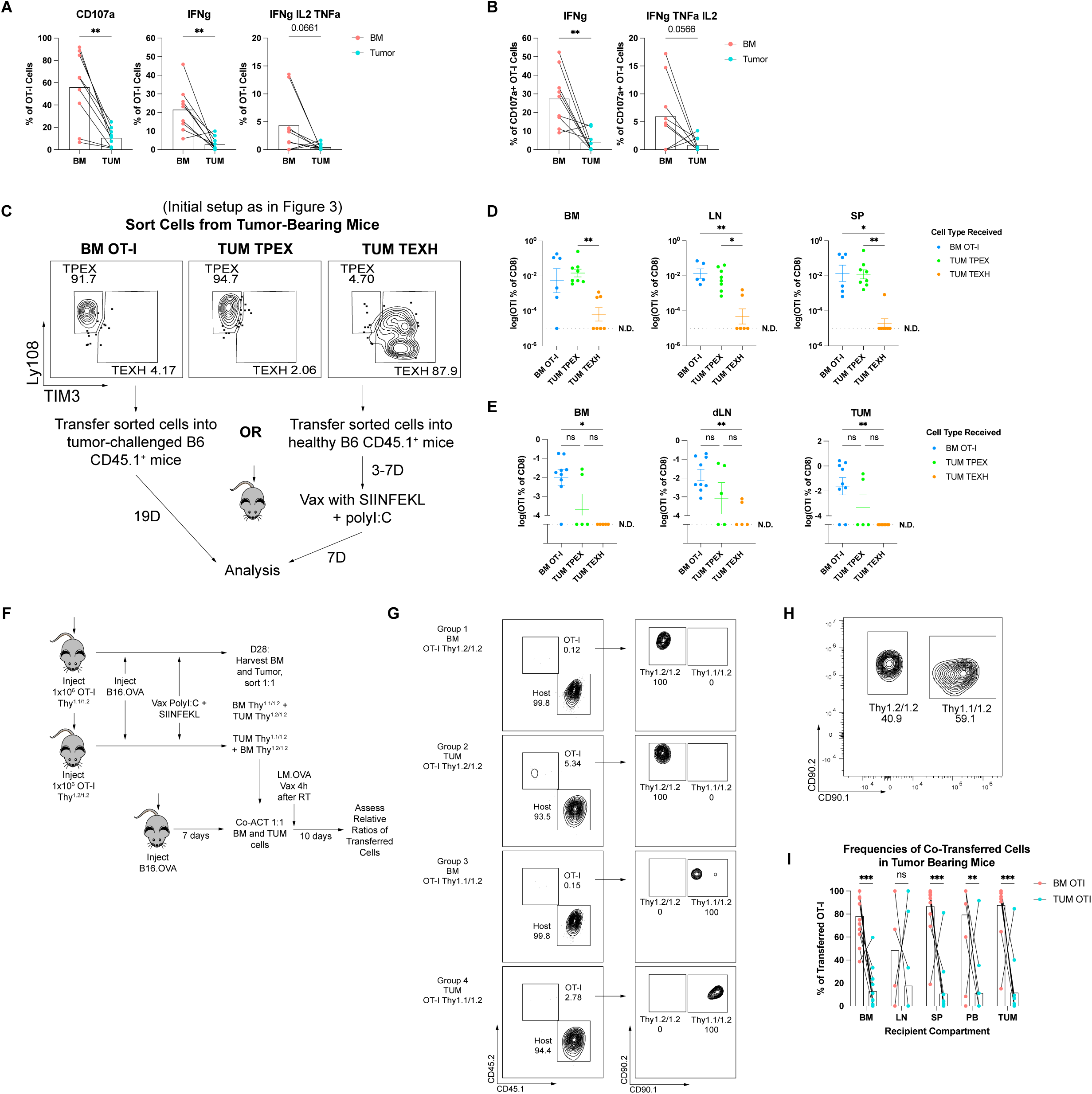
BM T cells exhibit greater persistence and tumor infiltration than TILs following adoptive transfer. **A:** CD107a expression, IFNγ production, and polyfunctional cytokine production by OT-I cells from BM and TUM following restimulation with PMA/Ionomycin. Summary of three replicate experiments, n=3 mice per experiment. Gated on CD45.2^+^ CD8^+^ T cells; analyzed with paired t test, **, p < 0.01. **B:** IFNγ and polyfunctional cytokine production by degranulating (CD107a^+^) OT-I cells from BM and TUM following restimulation with PMA/Ionomycin. Analyzed with paired t test, **, p < 0.01. **C:** Experimental overview of adoptive cell transfer to healthy and to tumor-bearing mice. BM OT-I (largely Ly108^+^), TUM TPEX (Ly108^+^TIM3^-^), or TUM TEXH (TIM3^+^Ly108^-^) cells were sorted from tumor-bearing mice for ACT to healthy recipients (which received vaccine) or mice challenged with tumor one day prior. **D:** Frequencies of BM OT-I cells, TUM TPEX and TUM TEXH in healthy recipients following vaccine challenge. Data from two replicate experiments, n=3-4 mice per group. Percent of OT-I values were log-transformed, and near-zero values were substituted for mice where 0% OT-I of CD8^+^ T cells were detected. Analyzed with Kruskal-Wallis test. *, p < 0.05; **, p < 0.01. **E:** Frequencies of BM OT-I cells, TUM TPEX and TUM TEXH in tumor-challenged recipients. Data from two replicate experiments, n=3-4 mice per group. Percent of OT-I values were log-transformed, and near-zero values were substituted for mice where 0% OT-I of CD8^+^ T cells were detected. Analyzed with Kruskal-Wallis test. *, p < 0.05; **, p < 0.01. **F:** Experimental schema for competitive co-adoptive cell transfer into tumor-challenged mice. **G:** Cell phenotype confirmation prior to sorting. OT-I cells are CD45.2^+^ and congenically marked with either CD90.2/CD90.2 (Thy^1.2/1.2^) or CD90.1/CD90.2 (Thy^1.1/1.2^). **H:** Representative data following 1:1 sort. For each pairing, BM OT-I cells were sorted first. Tumor OT-I cells with the opposite congenic marker were sorted into the same tube, with stopping gate set to match the number of cells derived from the BM. **I:** Frequencies of BM OT-I and TUM OT-I cells following adoptive co-transfer. Mice received 1500-3000 cells of each type per transfer. Data from three replicate experiments, n=3-6 mice per group.

Considering the significant phenotypic, transcriptional, and functional differences between BM T cells and TILs, we evaluated their activity *in vivo*. We assessed the response of BM OT-I cells (primarily Ly108^+^ TPEX) when compared to TUM TPEX (Ly108^+^ TIM3^-^) or TUM TEXH (TIM3^+^ Ly108^-^) to vaccine challenge as well as to tumor challenge (Fig 5C). Following adoptive transfer into healthy hosts with subsequent vaccine challenge (Fig 5D) or into tumor-challenged hosts (Fig 5E), BM OT-I cells and TUM TPEX were recovered at greater frequencies than TUM TEXH, which were rarely detected. While these experiments did not demonstrate significant differences in retrieval of BM OT-I and TUM TPEX as a percentage of all CD8^+^ T cells derived from recipients, the trends suggest that BM OT-I can be recovered more frequently than Ly108^+^ TUM TPEX, especially in tumor-challenged hosts (BM T cells were found in 7/9 recipients, 78% as compared to TUM TPEX found in 2/5 recipients, 40%). This indicates that TPEX-like cells derived from the BM exhibit distinct biology from TUM TPEX following adoptive transfer.

Lastly, in seeking to extend these observations, we compared BM T cells to TILs following co-adoptive transfer in tumor-bearing mice (Fig 5F). Congenically distinct OT-I cells were pooled from the BM or TUM of separate groups of tumor-bearing donors (Fig 5G), sorted at a 1:1 ratio (Fig 5H), and transferred into recipient mice challenged with B16.OVA. Four hours after transfer, recipients were challenged with Listeria monocytogenes expressing SIINFEKL peptide (LM-OVA). After ten days, we assessed the relative frequencies of adoptively transferred cells in tumor-bearing recipients. In this competitive transfer setting, BM T cells were found at greater frequencies than TILs in the BM and spleen, in the blood, and importantly within the tumor itself (Fig 5I). These data demonstrate that BM T cells persist upon transfer and infiltrate the tumor to a greater extent than TILs.

## Discussion

TILs have shown clinical efficacy in select solid cancers, including metastatic melanoma and lung cancer^6,49,50^. However, the successes of ACT in solid tumors are limited due to several unique challenges. These include a strongly immunosuppressive tumor microenvironment that can physically exclude cells and hamper T cell efficacy, the fact that not all tumor specimens yield enough T cells for expansion, and the laborious expansion process required to produce the final TIL product^7–9^. Additionally, as TILs are obtained from tumors, candidates for this therapy must possess a significant disease burden and be amenable to an elective surgical procedure to harvest the requisite T cell starting material. While TILs can generate clinically significant responses, these hurdles preclude the use of TILs in the context of early disease, in patients without harvestable tumors or in tumors with insufficient T cell infiltration.

Adoptively transferred cells must possess several key attributes: the capacity to traffic throughout the body and to the site of the tumor, effector functionality, the ability to proliferate and persist in the host through formation of memory, and the capacity to impart an anti-tumor effect *in vivo*. We and others have shown that BM T cells possess several of these attributes^10,16,17,31,32,34,46,51–54^. Specifically, the BM houses memory T cells specific for antigens from cancer and previous infections^16,25,28,30–33^. BM T cells possess enhanced reactivity to autologous tumor when compared to PBLs *in vitro* and *in vivo*^17,31,32,34^ and can induce a circulating memory response in patients which correlates with increased overall survival^54^. We sought to determine the extent to which BM cells differ from TILs as an alternative T cell source for ACT. We hypothesized that the inherent microenvironmental differences could drive distinct T cell phenotypes.

Profiling non-activated T cells allowed us to better understand the biological differences between BM T cells and TILs in their native state. We show that tumor-specific CD8^+^ T cells in the BM are phenotypically and transcriptionally more memory-like and less exhausted than TILs. Functionally, BM T cells are capable of degranulating and producing multiple effector cytokines. Importantly, BM T cells can infiltrate solid tumors and persist more than TILs following adoptive transfer. We hypothesize that the baseline enrichment for a tumor-specific memory T cell population in the BM is linked to the overall improved fitness demonstrated as compared to TILs following adoptive transfer. To our knowledge, this is the first study directly contrasting BM T cells against TILs and our findings underscore significant differences with important translational implications.

With an increasing knowledge of the heterogeneity that exists within T cell populations, approaches are being employed to select for subsets that will perform more optimally following adoptive transfer^55–58^. Murine studies have shown that central memory CD8^+^ T cells can give rise to effector T cells *in vivo* and sustain long-term immune memory following adoptive transfer to a greater extent than effector memory CD8^+^ T cells^56^. Additionally, manufacturing CAR-T cells from a selected subset of naïve or stem-like memory T cells (as opposed to bulk, non-selected T cells) enhances their efficacy and safety profile and yields a cell product with improved expansion and persistence *in vivo* after transfer, increased anti-tumor activity and decreased levels of cytokine release syndrome^55^. Several clinical studies have demonstrated that the T cell phenotype at baseline is predictive of clinical outcome following treatment with a cell product^31,55,56,59–62^. Specifically, treating patients with T cells that are more memory-like or stem-like prior to expansion correlates with: sustained remissions ^61^, improved overall response rates ^31^, and increased overall survival ^54^. Collectively, these data indicate that the baseline cell phenotype plays a critical role in determining T cell persistence and functionality following adoptive transfer.

The quality of the initial T cell population is a critical factor linked to treatment outcomes following ACT. Regardless of whether they are progenitor-like, effector-like, or terminally exhausted, tumor-specific TILs are dysfunctional. They produce fewer effector cytokines and fail to successfully form memory, to mount responses to cognate antigen upon re-challenge, and to persist *in vivo* ^63–66^. Moreover, truly exhausted T cells are resistant to reprogramming^64,66,67^. As it is a reservoir of memory CD8^+^ T cells with progenitor-like attributes that are less exhausted than TILs, the BM could be a viable cell source for ACT in solid malignancies. Utilizing the BM instead of TILs enables the development of a T cell product with T cells that are less dysfunctional, yet still tumor-specific. Additionally, considering that tumor-specific T cells are present in the BM in non-metastatic disease, this approach could prove viable in earlier stages of disease where harvesting tumor is not feasible.

At present, the factors facilitating memory enrichment within the BM niche remain to be fully elucidated and the role of the BM in anti-tumor immunity is incompletely understood. It is unclear whether endogenous tumor-specific BM T cells originate in the BM (and thus, are primed there), or if they are primed elsewhere and then traffic to and lodge within the BM. Published findings support either position^16–18,24,51,68–71^, and our model does not address this question.

Our work highlights the marked differences in T cell exhaustion states between BM T cells and TILs. Notably, in this model tumor is not present in the BM and thus these significant differences in T cell function could simply be explained by the presence or absence of tumor. Earlier studies in myeloma demonstrated that, despite the presence of a higher disease burden in the BM, BM T cells exhibit less exhaustion compared to T cells from the peripheral blood^72^. These data suggest that the BM microenvironment is capable of maintaining a less exhausted T cell phenotype, independent of the presence of tumor burden

The data shows that the BM houses tumor-specific memory-like CD8^+^ T cells in solid tumors and corroborates data from human studies showing greater tumor specificity in the BM of solid tumor patients with non-metastatic disease compared to the blood. Tumor-specific, BM-derived T cells are not phenotypically or transcriptionally exhausted and persist longer than TILs in tumor-bearing hosts. These data suggest that approaches utilizing BM T cells for adoptive cell therapy against solid cancers warrant further investigation.

## Methods

### Basic B16.OVA Model Setup

B16.OVA cells were a gift from the lab of Jonathan Powell. B6 CD45.1^+^ mice were implanted subcutaneously with 250,000 B16.OVA tumor cells. Cells were cultured in complete RPMI medium under g418 selection (400μg/mL). At select tumor timepoints, mice were sacrificed, tissues were processed as described below, and cells were immunophenotyped using flow cytometry.

### Adapted B16.OVA Model Setup

Spleens were harvested from 6-10-week-old C57Bl/6 OT-I transgenic mice as a source of naïve OT-I cells. Single cell suspensions of splenocytes were generated by disaggregation through a strainer, followed by red blood cell lysis. CD8^+^ T cells were enriched using the StemCell EasySep CD8^+^ T cell kit (#19853). C57Bl/6 CD45.1^+^ mice were injected with 1×10^6^ naïve OT-I cells. One day later, the mice received 250,000 B16.OVA tumor cells subcutaneously on the right flank (D0). Three days following tumor injection, mice received subcutaneous vaccine comprised of 50μg polyI:C and 10μg SIINFEKL peptide in the opposite flank. Tumors were allowed to grow out until analysis endpoints specified for individual experiments.

### Tissue Processing

The standard tissue processing protocol was as follows. <u>Bone Marrow</u>: Bone marrow was flushed from the femur and tibia with a 27.5-gauge needle with PBS onto a pre-wetted 70uM strainer. Cells were pressed through a strainer and washed thoroughly with PBS. Samples underwent red blood cell lysis using ACK lysis buffer (Quality Biological #118-156-101), and were washed with PBS. Bone marrow was enriched for T cells with either CD3 or CD8 enrichment kits from StemCell (#19851 or #19853). In cases where absolute purity was not necessary (e.g. downstream sorting applications or flow cytometry) enrichment was deliberately imperfect to preserve reagents. In instances when cells were intended for adoptive transfer, manufacturer protocol was followed, yielding >90% T cell purity. <u>Draining Lymph Node</u>: The dLN was harvested from tumor-bearing mice, minced on a pre-wetted 70uM strainer, pressed through with a plunger and rinsed with PBS. <u>Spleen</u>: Spleens were harvested and pressed through a pre-wetted 70uM strainer and washed with PBS. Following this wash, samples underwent red blood cell lysis using ACK lysis buffer and were rinsed with PBS. <u>Tumors</u>: Tumors were minced and incubated in cRPMI with 10IU/mL (20μg/mL) DNAse and 200IU/mL type IV collagenase for 45 minutes at 37C under agitation. The tumors were then pressed through a 70uM strainer and repeatedly rinsed with PBS. Lymphocytes were enriched from tumors using a gradient of Ficoll-Paque PREMIUM (1.084g/mL (Cytiva #17544602).

### Polychromatic Flow Cytometry Staining and Panel Development

For all incubation steps, samples were protected from light. Samples for staining were washed 2x with PBS and stained with Zombie NIR live/dead discriminant dye (2000x) for 20 minutes at RT. Following viability staining, samples were washed with FACS buffer (4% FBS in PBS). When applicable, OVA-Tetramer stain was applied at room temperature, followed by 1x wash. Chemokine stain was applied and samples were incubated for 30 minutes at 37C. Samples were washed 1x with FACS buffer. Surface stain was applied to samples, which remained at room temperature for 30 minutes. For experiments involving sorting, samples were resuspended in 1% FBS. When applicable, samples were either fixed with the Cytofix/Cytoperm Plus Kit (BD #555028) (for cytokine staining) or the True-Nuclear Transcription Factor Buffer Set (Biolegend #424401) (for intranuclear staining) for 30 minutes at room temperature. Antibody master mixes for intracellular or intranuclear stain were resuspended in the respective perm/wash buffer and applied for 30 minutes. Prior to data collection, samples were washed with and resuspended in FACS buffer.

Data were collected on a Cytek Northern Lights spectral flow cytometer equipped with 3-lasers (405nm, 488nm, 640nm). A full list of the antibodies used is provided in Supplementary Table 1. Flow cytometry controls were generated from splenic murine T cells and/or beads for spectral unmixing, choosing the optimal control per fluorochrome after testing both means. Antibodies were previously titrated to the optimal concentration.

### Cell Sorting

For adoptive transfer experiments and experiments with downstream *in vitro* applications, cells were sorted into 4% FBS buffered solution on a BD FACSAria Fusion Sorter. Sorted cells were washed with PBS or cRPMI as appropriate at 400xg for 8 minutes prior to further use. For RNA Sequencing, cells were sorted directly into TrizolLS, snap frozen, and kept at -80C until further processing.

### Mice

Wild-type C57Bl/6 (JAX #000664), CD45.1^+^ C57Bl/6 (B6.SJL-*Ptprc^a^ Pepc^b^*/BoyJ, JAX #002014), transgenic OT-I (C57BL/6-Tg(TcraTcrb)1100Mjb/J, JAX #003831), and B6 Thy1.1 (B6.PL-*Thy1^a^*/CyJ, JAX #000406) mice were obtained from Jackson labs. Thy1.1/1.2 expressing OT-I mice were bred by crossing B6 OT-I with B6 Thy1.1/1.1 mice; heterozygosity for the Thy1.1/1.2 allele as well as Tet^+^ nature of OT-I cells were confirmed via flow cytometry. Mice aged 4-12 weeks were used for all experiments. Mice used as donors were male and female; mice used as recipients were female. All animal studies were approved by and conducted in accordance the Johns Hopkins University Institutional Animal Care and Use Committee.

### *Ex Vivo* Re-stimulation Assay

The model was set up as referenced above (adapted B16.OVA model setup). Tumors were established for 28-32 days. BM underwent CD3 enrichment via StemCell Mouse T cell Isolation Kits, TUM were processed with Ficoll, and SP and dLN were not enriched. Following processing, samples were plated in a 96-well, round-bottom plate and resuspended in complete RPMI media containing 500x dilution of Cell Stimulation Cocktail (PMA / Ionomycin, eBioscience 00-4970-93), 500x of protein secretion inhibitor containing GolgiStop and GolgiPlug (eBioscience #00-4980-93), and 2μL per well of AF488-conjugated anti-mouse CD107a. Samples were incubated at 37C for 3.5-4 hours.

### Adoptive Transfer of Sorted BM OT-I, TUM TPEX, or TUM TEXH

C57Bl/6 CD45.1^+^ donor mice underwent adapted model experiment setup. At D27 following tumor inoculation, the following subsets were sorted from pooled samples: OT-I cells (CD45.2^+^) derived from the BM (>90% Ly108^+^ TPEX), TUM TPEX (Ly108^+^TIM3^-^), TUM TEXH (TIM3^+^Ly108^-^). <u>Transfer into healthy recipients</u>: Sorted cells were adoptively transferred via tail-vein injection into healthy C57Bl/6 CD45.1^+^ mice (3-5 recipient mice per group). 3-7 days later, recipient mice received subcutaneous polyI:C and SIINFEKL vaccine on the flank. 7 days following vaccination, mice were sacrificed, BM, LN and SP were harvested and processed as usual, and samples were stained for flow cytometry to assess frequency of transferred cells. <u>Transfer into tumor-challenged recipients</u>: Sorted cells were adoptively transferred via tail-vein injection into C57Bl/6 CD45.1^+^ mice which had received 125,000 B16.OVA cells subcutaneously the previous day. Tumors were monitored every 3-4 days. After tumor outgrowth for 18-19 days, mice were sacrificed, BM, dLN and TUM were harvested, processed as usual, and samples were stained for flow cytometry to assess frequency of transferred cells.

For both sets of experiments, frequencies of CD45.2^+^ OT-I cells within CD8^+^ T cells were log-transformed and analyzed using the Kruskal Wallis test.

### Co-Adoptive Transfer of BM and TUM Cells

<u>Donor mice</u>: Separate cohorts of C57Bl/6 CD45.1^+^ mice were injected with 1×10^6^ OT-I cells expressing different congenic markers: Thy1.1/Thy1.2 (CD45.2^+^CD90.1/CD90.2^+^) or Thy1.2 homozygous (CD45.2^+^ CD90.2/CD90.2^+^). Mice received 250,000 B16.OVA cells subcutaneously on day 0 and were challenged with polyI:C + SIINFEKL vaccine 3 days later. <u>Cell harvest, sorting and adoptive transfer</u>: On D28, donor mice were sacrificed, bone marrow and tumor cells T cells were harvested, processed, and pooled into 4 groups: BM OT-I CD90.2/CD90.2; BM OT-I CD90.1/CD90.2; TUM OT-I CD90.2/CD90.2; TUM OT-I CD90.1/CD90.2. Cells were stained with fluorochrome-conjugated antibodies to discriminate their phenotype and prepared for cell sorting. BM and TUM-derived OT-I cells with opposite congenic labels were sorted at a 1:1 ratio and adoptively transferred into tumor-bearing C57Bl/6 CD45.1^+^ recipients challenged with B16.OVA 7 days prior. Recipient mice received between 2000 and 3000 total OT-I cells (1000-1500 from BM and 1000-1500 from TUM). 4 hours after adoptive transfer, recipients were challenged with 2 x 10^6^ CFU of OVA-expressing Listeria monocytogenes (DactA, Din1B) (LM-OVA) (gift from Aduro Biotech). <u>Experiment</u> <u>endpoint and flow cytometry</u>: Ten days after co-transfer, recipient mice were sacrificed and BM, dLN, SP and TUM were collected and analyzed to assess relative ratios of BM-and TUM-derived OT-I cells.

### RNA Sequencing

C57Bl/6 CD45.1^+^ mice underwent the adapted model setup. At D7 following tumor injection, bone marrow was harvested from 5 mice, processed, and prepared for sorting. Tumors were just barely visible under the skin at this time. TPEX (Ly108^+^) and TEFF (Ly108^+^ CX3CR1^+^) subsets were sorted from BM samples, snap frozen, and stored at - 80C. At D28, bone marrow and tumors were isolated from 5 mice, processed and prepared for sorting. TPEX (Ly108^+^) and TEFF (Ly108^+^ CX3CR1^+^) cell subsets were sorted from bone marrow and tumor samples, along with TTERM cells (Ly108^-^CX3CR1^-^) from tumors. Samples were snap frozen and stored at -80C.

### Ultra-low input RNA-seq

Following collection of samples from both timepoints, RNA Extraction, samples QC, library preparations, sequencing reactions, and initial bioinformatic analysis were conducted at GENEWIZ/Azenta Life Sciences LLC. (South Plainfield, NJ, USA).

#### RNA Extractions and sample QC

Total RNA was extracted from cells following the Trizol Reagent User Guide (Thermo Fisher Scientific). Extracted RNA samples were quantified using Qubit 2.0 Fluorometer (Life Technologies, Carlsbad, CA, USA) and RNA integrity was checked using Agilent TapeStation 4200 (Agilent Technologies, Palo Alto, CA, USA).

#### Library preparation and Sequencing

The SMARTSeq HT Ultra Low Input Kit was used for full-length cDNA synthesis and amplification (Clontech, Mountain View, CA), and Illumina Nextera XT library was used for sequencing library preparation. Briefly, cDNA was fragmented, and adaptor was added using Transposase, followed by limited-cycle PCR to enrich and add index to the cDNA fragments. Sequencing libraries were validated using the Agilent TapeStation and quantified by using Qubit Fluorometer as well as by quantitative PCR (KAPA Biosystems, Wilmington, MA, USA). The sequencing libraries were multiplexed and clustered on a flowcell. After clustering, the flowcell was loaded on the Illumina HiSeq instrument according to manufacturer’s instructions. The samples were sequenced using a 2×150 Paired End (PE) configuration.

#### Bioinformatic Analysis

Image analysis and base calling were conducted by the HiSeq Control Software (HCS). Raw sequence data (.bcl files) generated from Illumina HiSeq was converted into fastq files and de-multiplexed using Illumina’s bcl2fastq 2.17 Software. One mismatch was allowed for index sequence identification. After investigating the quality of the raw data, sequence reads were trimmed to remove possible adapter sequences and nucleotides with poor quality using Trimmomatic v.0.36. The trimmed reads were mapped to the Mus musculus reference genome available on ENSEMBL using the STAR aligner v.2.5.2b. The STAR aligner is a splice aligner that detects splice junctions and incorporates them to help align the entire read sequences. BAM files were generated as a result of this step. Unique gene hit counts were calculated by using feature Counts from the Subread package v.1.5.2. Only unique reads that fell within exon regions were counted. After extraction of gene hit counts, the gene hit counts table was used for downstream differential expression analysis.

### Additional Bioinformatic Analysis

Upon receipt of the preliminary analysis data package, counts data were extracted, combined, and put into a custom pipeline in RStudio for scaling, hierarchical clustering, identification of differentially expressed genes (DEGs), *k means* clustering, as well as gene set enrichment analysis (GSEA).

Briefly, using the R/Bioconductor *edgeR* (v3.40.2) package^75^, the M-value trimmed mean method (TMM) was employed to normalize samples by library size, filter out genes with low counts, and to compute the gene expression level in log2 counts per million^76,77^. Differential expression (DE) analysis and linear modeling were performed with *limma* (v.3.54.2) to compare expression levels between tissue (BM vs. TUM), subsets between tissues (e.g BM TPEX vs. TUM TPEX and BM TEFF vs. TUM TEFF), or subsets within a tissue (e.g. BM TPEX vs. TEFF).

For gene set enrichment analysis (GSEA), genes were ranked according to the t statistic obtained from the DE analysis. The R package *fgsea* (v1.24.0) ^78–80^, which employs a multilevel split Monte Carlo approach ^78^, was then used to estimate relative p-values and enrichment score of gene sets obtained from the Molecular Signature Database (MSigDB). For all analyses, multiple comparisons testing corrections were performed with the Benjamini-Hochberg method. The *tidyverse* (v2.0) packages were used for data handling and visualization ^81^.

Data will be deposited in the GEO database and accession number will be available upon reviewer request.

### Analysis of Flow Cytometry Data

Data were analyzed using a combination of manual gating and computational methods. Manual gating was performed using FlowJo analysis software, and numerical data were statistically analyzed via GraphPad Prism.

For computational analyses, populations of interest for a given analysis (e.g. OT-I, CD8, TPEX, TEFF or TTERM) were manually gated, cleaned to remove dead cells, doublets, and aggregates, and exported from FlowJo for further analysis in R (version 4.2.2) using a custom-made script comprised of Bioconductor libraries and R packages which was developed by L. Biavati. Briefly, data were analyzed using the FlowSOM algorithm for unsupervised clustering and then visualized with Uniform Manifold Approximation and Projection (UMAP). Differential discovery analyses were performed using the *diffcyt* framework and the CATALYST workflow^83^.

### Code and data availability

Materials generated by this study are available upon request, custom-made R scripts will be made available via Github.

### Statistical Analysis

All data are presented as the mean (for paired analyses) or mean +/- SEM. Statistical analyses were performed using GraphPad Prism 6 with tests appropriate for experimental setup, as annotated in the figure legends. Comparisons of a single variable measured in a sample at two different sites (e.g. tissues from an individual mouse) were performed via a paired, two-tailed Student’s t-test. Comparisons of one variable across multiple conditions were conducted using a one-way ANOVA with Tukey’s correction for multiple comparisons. Comparisons of one variable across multiple conditions with non-Gaussian distribution were performed using the Kruskal-Wallis test with Dunn’s correction for multiple comparisons. Significance levels are reported as follows: ****, p<0.0001; ***, p<0.001; **, p<0.01; *,p<0.05.

