## Supplementary figures and images for "Tumor-Specific CD8^+^ T Cells from the Bone Marrow Resist Exhaustion and Exhibit Increased Persistence in Tumor-Bearing Hosts as Compared to Tumor Infiltrating Lymphocytes"

### Supplemental_materials_manuscript_.pdf

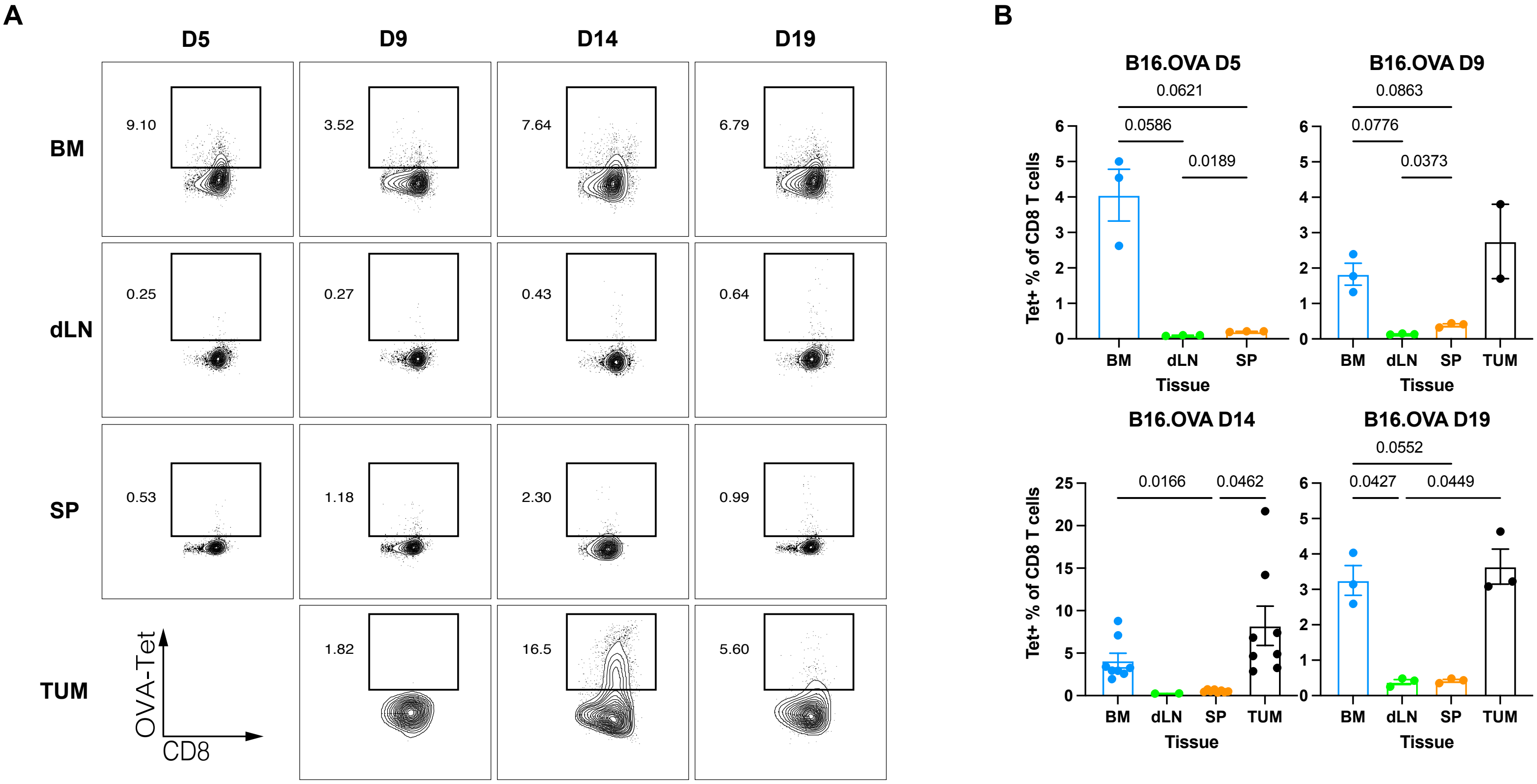

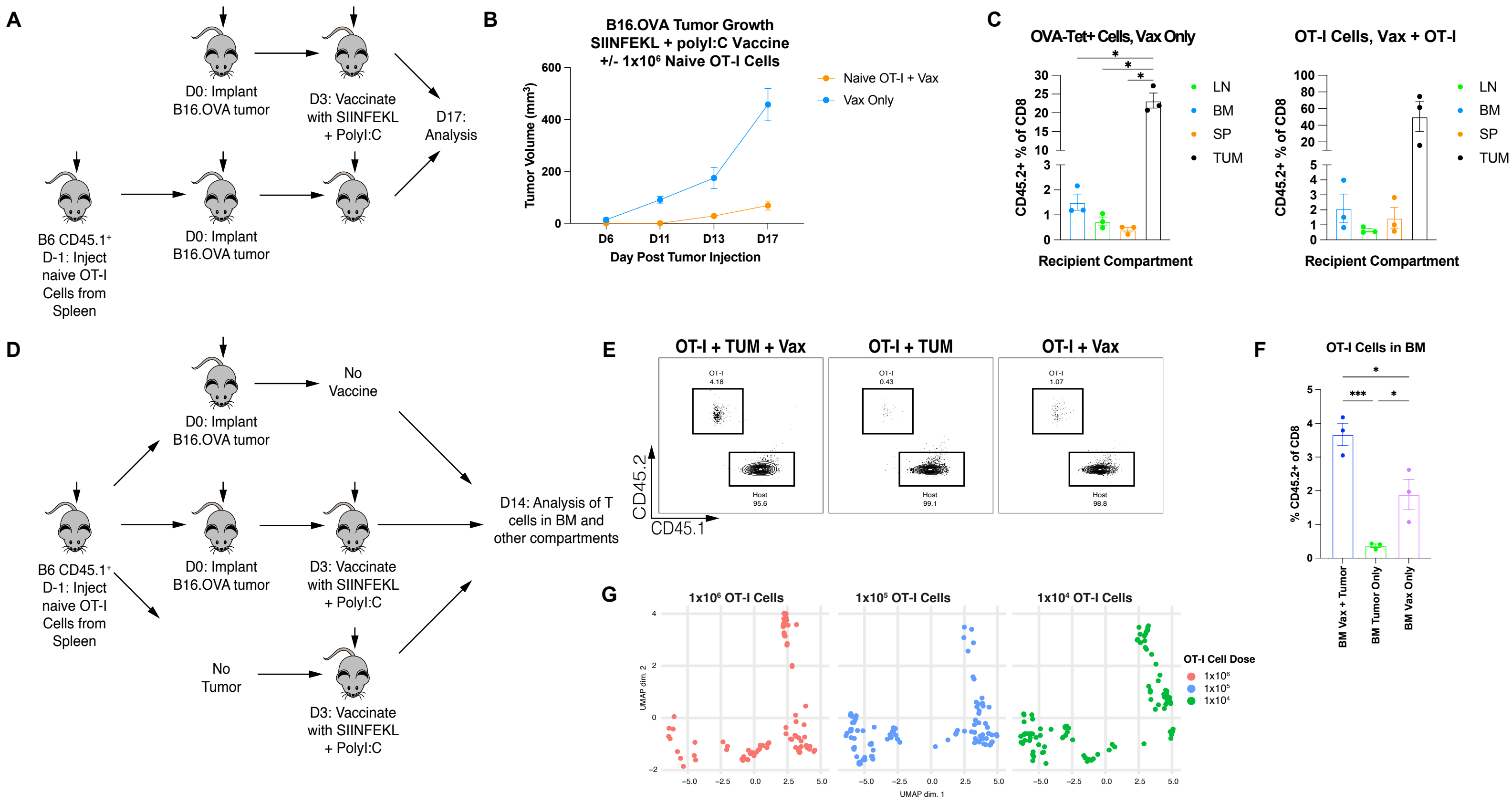

#### Supp Fig 3

A

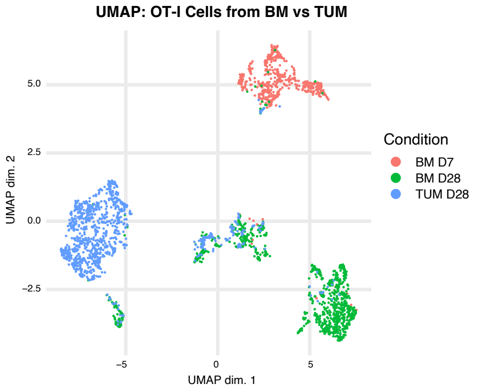


B

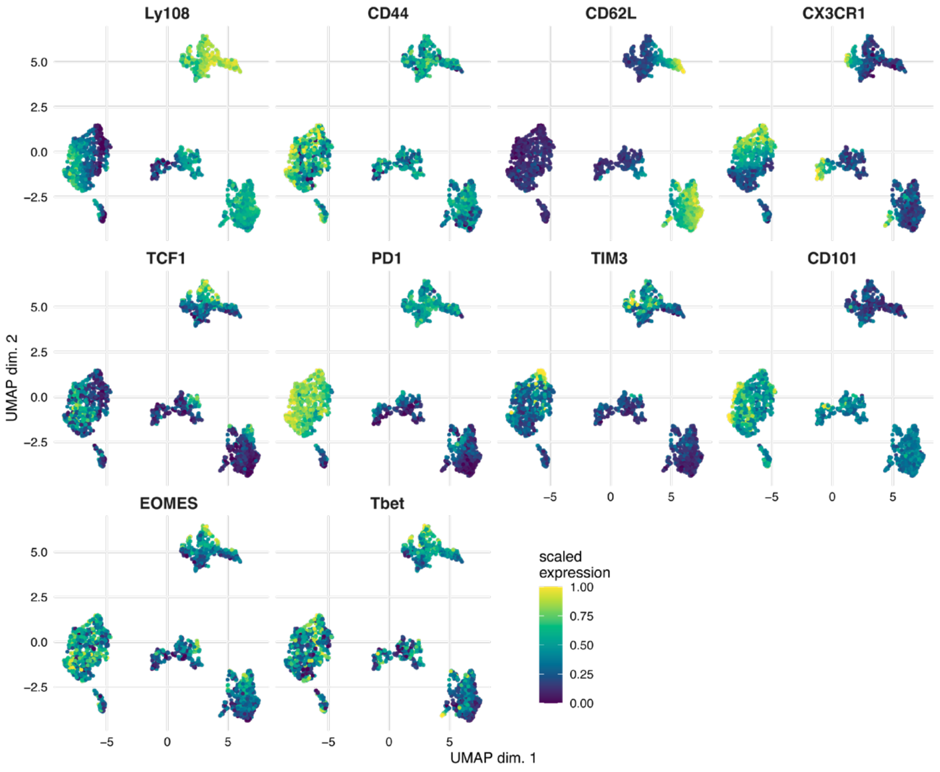


C

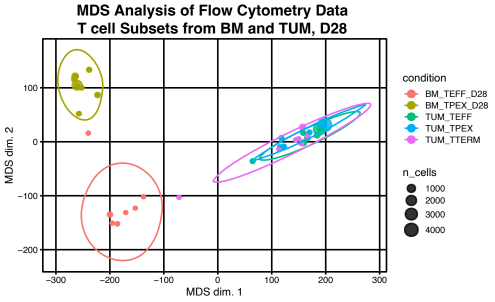

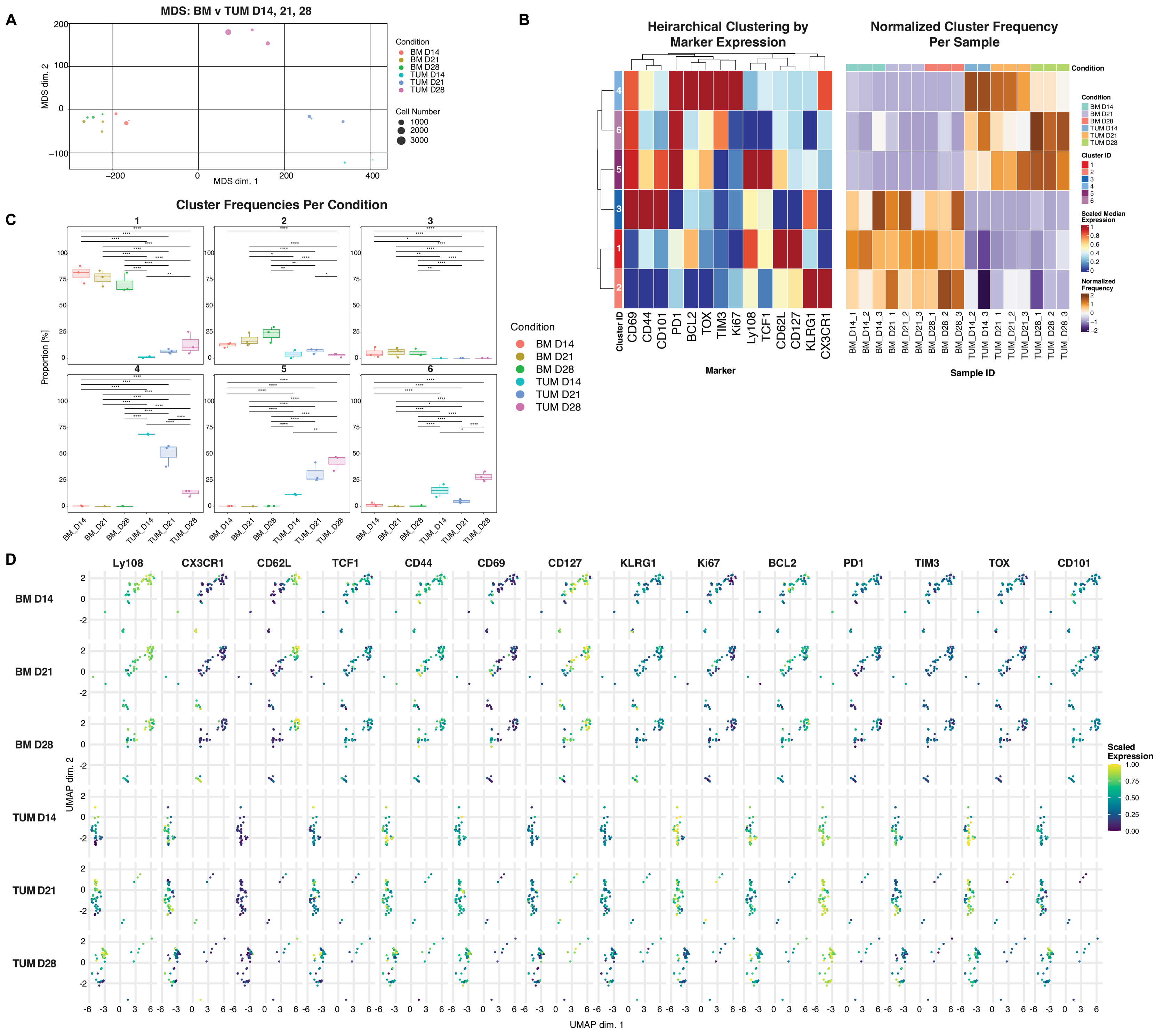

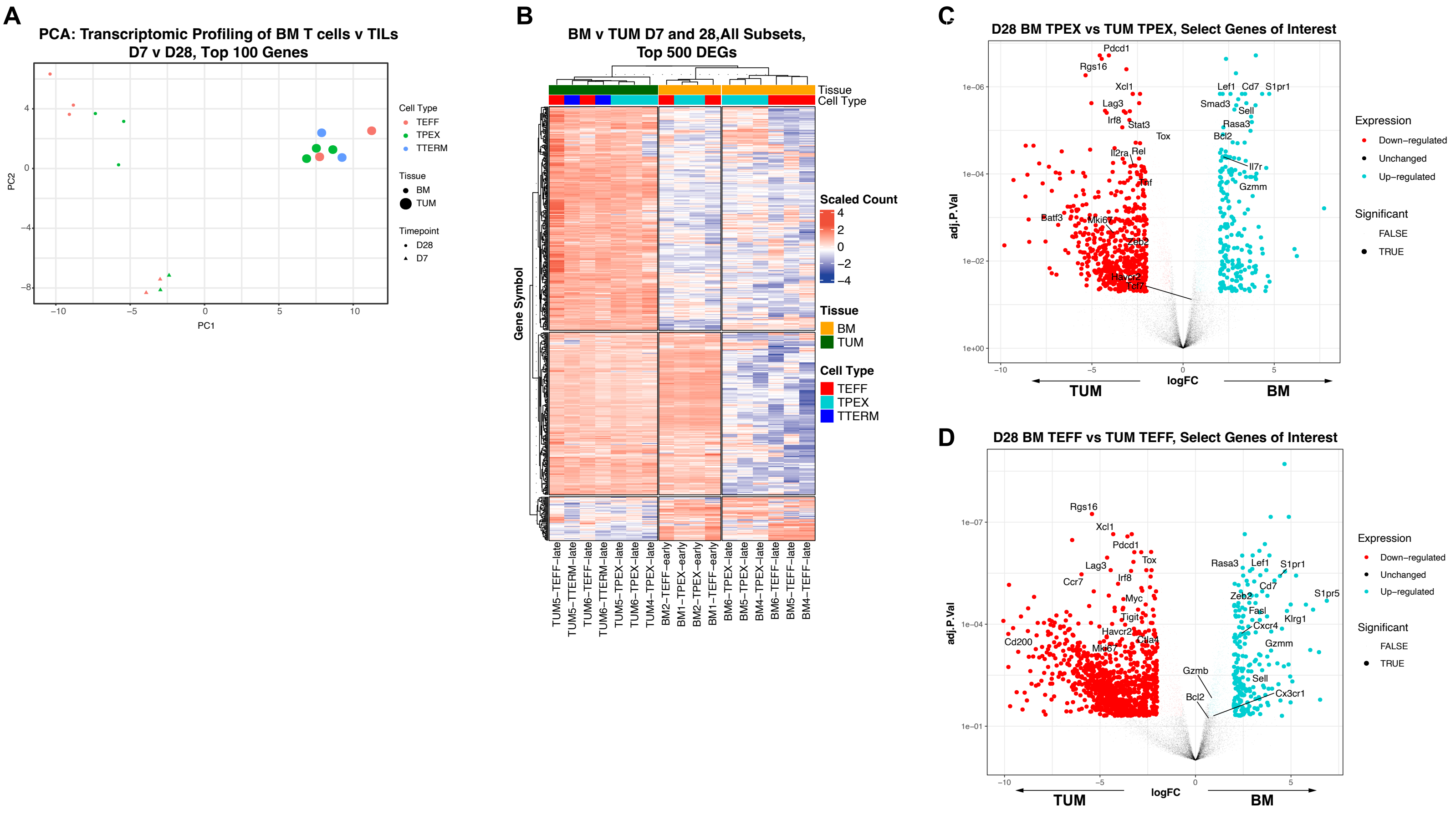

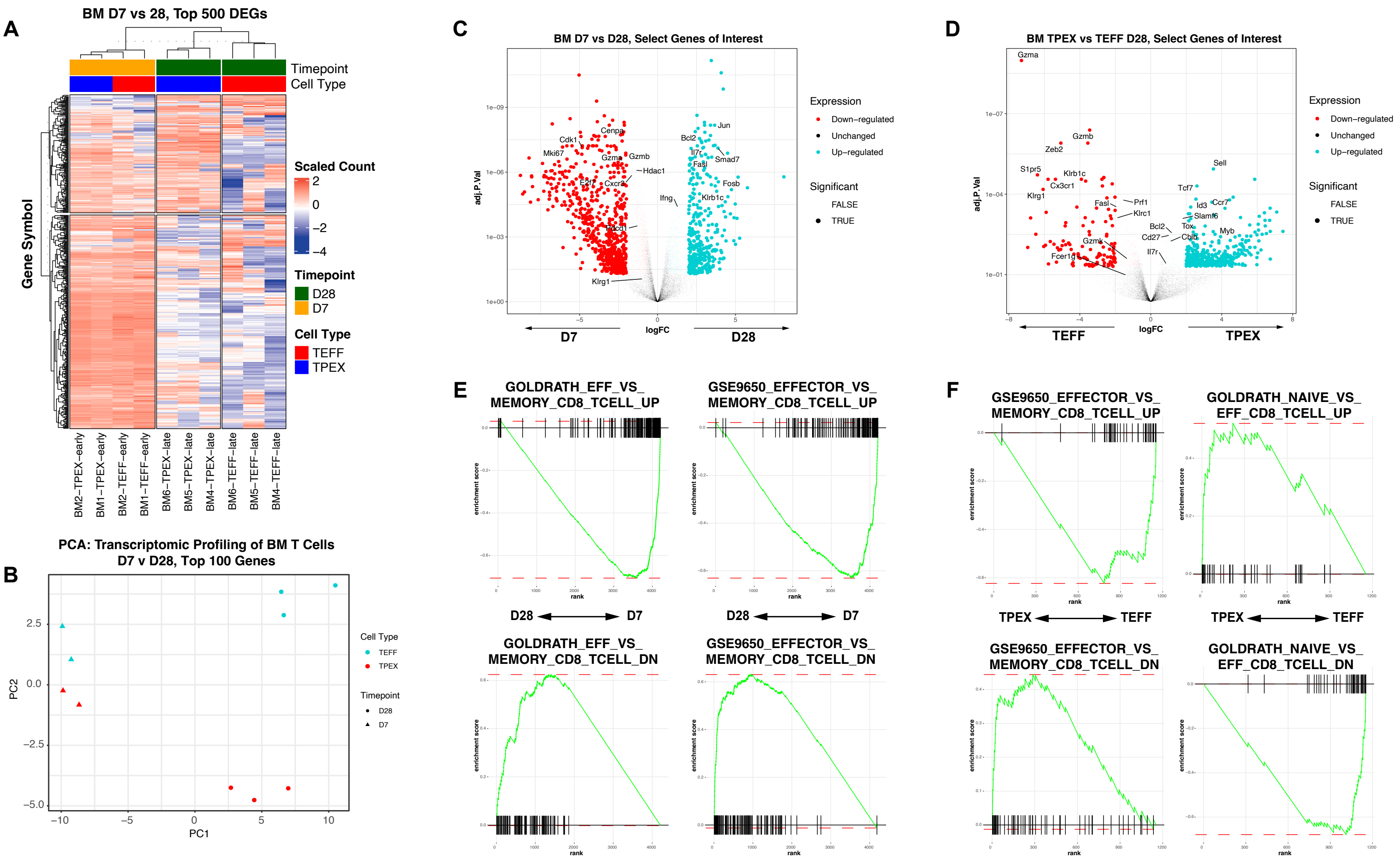

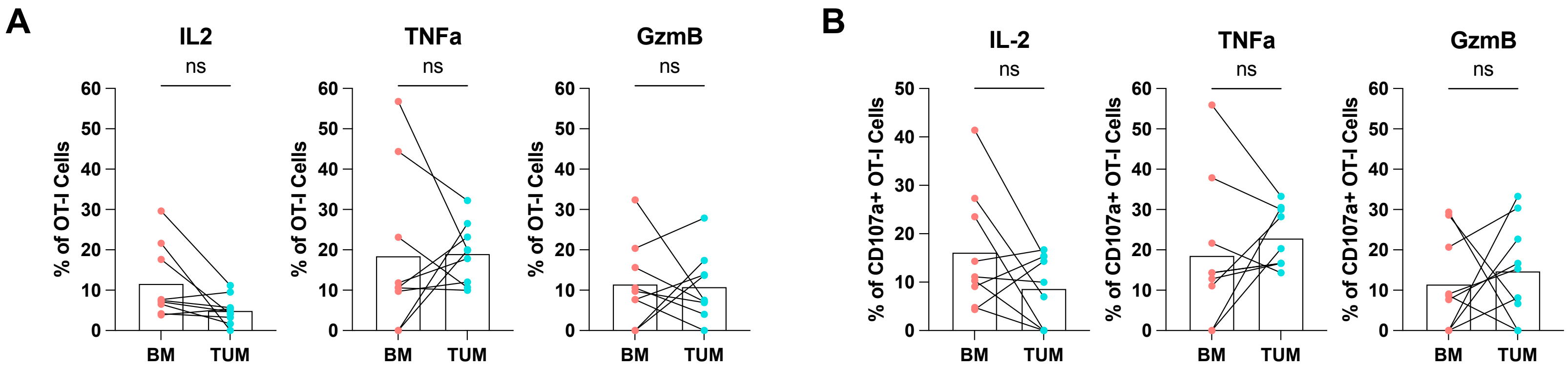
