## Supplemental Figure Legends for "Tumor-Specific CD8^+^ T Cells from the Bone Marrow Resist Exhaustion and Exhibit Increased Persistence in Tumor-Bearing Hosts as Compared to Tumor Infiltrating Lymphocytes": Figure Legends_SupplementalFigures.pdf

### Supplemental Data

**Supplementary Figure 1: The BM maintains a reservoir of OVA-Tet$^+$ CD8$^+$ T cells during B16.OVA tumor progression.**
**A:** Flow cytometry representative data from timecourse experiment. Mice received subcutaneous B16.OVA tumor and were sacrificed for analysis at different timepoints.
**B:** Quantification of Tet$^+$ CD8$^+$ T cells in different tissues at different timepoints following tumor injection. n=2-5 mice per group, summary of two different experiments. RM one-way ANOVA $F(2,6) = 28.69$, $p=0.0008$. p values for multiple comparisons testing as indicated.

**Supplementary Figure 2: Validation of adapted B16.OVA model with naïve OT-I cells.**
**A:** Workflow for comparison of endogenous vs. adoptively transferred T cells in the B16.OVA model.
**B:** Tumor growth in mice receiving either B16.OVA plus vaccine only or mice which received ACT of $1x10^6$ naïve OT-I cells prior to tumor injection.
**C:** Frequencies of OVA-Tet$^+$ CD8$^+$ T cells in different organs following vaccination with SIINFEKL + polyI:C (RM one-way ANOVA $F(1.047, 2.094) = 111.2$, $p = 0.0076$) or frequencies of OT-I cells in different organs following vaccination (not significant in this example, RM one-way ANOVA $F(1.001, 2.001) = 7.219$, $p = 0.1150$). Asterisks indicate p values for multiple comparisons testing. *, $p < 0.05$.
**D**: Workflow to compare the effect of vaccine, tumor, or both tumor and vaccine on yield of OVA-specific OT-I cells from the BM.
**E:** Representative data from the bone marrow of mice receiving different treatments, n=3 mice per treatment group.
**F:** Frequencies of OT-I cells in the BM per indicated condition. Ordinary one-way ANOVA $F(2,6) = 26.12$, $p = 0.0011$. Asterisks indicate p values for multiple comparisons testing. *, $p < 0.05$; ***, $p < 0.001$.
**G:** UMAP of flow cytometry data profiling OT-I cells derived from the BM and TUM of mice which received different doses of naïve OT-I cells prior to injection of B16.OVA tumor in the adapted model workflow. Analysis examined key cell state markers via high-dimensional analysis (CD62L, CD44, PD1, OVA-Tet, CX3CR1, CD69, Tbet, TIM3, TCF1, EOMES, CD101).

**Supplementary Figure 3: At D7 and D28, adoptively transferred, monoclonal OT-I cells derived from the BM and TUM exhibit different phenotypes.**
**A:** UMAP analysis of OT-I cells from D7 BM, D28 BM, and D28 TUM (early and late timepoints).
**B:** Projection of individual markers onto the UMAP in A.
**C:** MDS analysis of flow cytometry phenotyping data from D28 BM and TUM, broken down by Ly108/CX3CR1-defined subset (TPEX, TEFF, TTERM).

**Supplementary Figure 4: Key phenotypic changes that are observed in T cells from the BM or the TUM manifest over time.**

**A:** MDS analysis of flow cytometry phenotyping data from BM and TUM on D14, D21 and D28 of tumor progression.
**B:** Marker expression by individual cluster from UMAP analysis.
**C:** Individual cluster frequencies from unsupervised clustering of data in UMAP analysis.
**D:** Individual phenotypic marker expression on cells derived from BM or TUM, separated by tumor timepoint.

**Supplementary Figure 5: Additional transcriptomic profiling of T cells from BM and TUM at early and late tumor timepoint.**
**A:** PCA of top 100 differentially expressed genes from TPEX, TEFF and TTERM subsets from BM and TUM at D7 and D28 of tumor progression.
**B:** Top 500 differentially expressed genes in T cell subsets derived from BM or TUM at D7 and D28, clustered using *k*-means.
**C:** Volcano plots of BM TPEX vs. TUM TPEX with select genes highlighted. Upregulated = upregulated in the BM; downregulated = downregulated in the BM and enriched in the tumor.
**D:** Volcano plots of BM TEFF vs. TUM TEFF with select genes highlighted. Upregulated = upregulated in the BM; downregulated = downregulated in the BM and enriched in the tumor.

**Supp Figure 6: Tumor-specific T cells found in the BM undergo marked changes during tumor progression**.
**A:** Heatmap of top 500 differentially expressed genes between OT-I cells derived from BM at early (D7) and late (D28) tumor timepoint.
**B:** PCA analysis of top 100 DEGs in BM OT-I cells.
**C:** Volcano plot showing select genes of interest upregulated in BM OT-I at D28 versus D7.
**D:** Volcano plot showing select genes of interest upregulated in BM OT-I TPEX versus TEFF at tumor D28.
**E:** GSEA comparing all BM OT-I cells with TUM-derived OT-I cells at tumor D28.
**F:** GSEA comparing BM TPEX vs. TEFF cells at tumor D28.

**Supp Figure 7: BM T cells and TILs exhibit similar production of select effector cytokines upon restimulation.**
**A:** Single effector cytokine production by OT-I cells from BM and TUM following restimulation with PMA/Ionomycin. of three replicate experiments, n=3 mice per experiment. Gated on CD45.2$^+$ CD8$^+$ T cells; analyzed with paired t test, **, $p < 0.01$.
**B:** Single effector cytokine production by degranulating (CD107a$^+$) OT-I cells from BM and TUM following restimulation. **, $p < 0.01$.
